# Clinical stringent response activation promotes conjugal transfer of staphylococcal resistance plasmids

**DOI:** 10.1101/2025.11.12.688057

**Authors:** Ashley T. Deventer, Ava Sutherland, Daria Biernacka, Paul R. Johnston, Claire E. Stevens, Anna-Karina Kaczorowska, Alisdair B. Boraston, Joanne K. Hobbs

## Abstract

Conjugative transfer of plasmids represents a major route through which antibiotic resistance genes are spread. In the case of the prevalent and deadly pathogen *Staphylococcus aureus*, >90% of clinical isolates carry at least one plasmid. While plasmid-encoded mechanisms (*e.g.* plasmid copy number) can influence conjugation frequency, host factors and environmental stimuli can also affect transmission. In particular, stress responses like the stringent response have been associated with increased movement of mobile genetic elements. We have previously shown that clinical mutations in the stringent response controller, Rel, lead to elevated levels of the alarmones (p)ppGpp and antibiotic tolerance in *S. aureus*. Here, we report that stringent response activation in these strains promotes the conjugal transfer of diverse staphylococcal resistance plasmids. We observed that clinical Rel mutations promote donation, but not receipt, of plasmids from the three families of staphylococcal plasmid and a mobilisable plasmid. This increased conjugation frequency could also be induced by chemical induction of the stringent response by mupirocin. Intriguingly, detailed experimental analysis revealed that the effect of elevated (p)ppGpp on plasmid donation was not due to CodY derepression, SOS response induction, increased plasmid copy number or increased expression of conjugation machinery genes. Further, transcriptomic analysis failed to identify any other putative plasmid- or host-derived mechanisms to explain this observation. Further investigations are required to explore the mechanistic link between the stringent response and conjugation, given the pervasive transcriptional and post-translational effects of (p)ppGpp. Overall, the association between Rel mutation and increased plasmid donation is alarming, especially as Rel mutations are being increasingly identified among clinical isolates.

## INTRODUCTION

Horizontal gene transfer and mobile genetic elements (MGEs), like plasmids, are the main drivers behind the spread of antibiotic resistance (1–3). Bacterial conjugation describes the process whereby genetic material is passed between two cells *via* direct contact. Conjugative plasmids are extra-chromosomal DNA elements that encode for all of the machinery necessary for their transfer, including mating pore proteins and a suite of other proteins required for DNA processing, replication and recruitment (4). These plasmids also frequently encode antibiotic resistance genes, sometimes as many as 14 (5). Conjugative plasmids are also capable of mobilising non-conjugative plasmids when they reside in the same cell; estimates suggest that at least 80% of non-conjugative plasmids found in *Staphylococcus aureus* can be mobilised (4). Therefore, conjugation is an important driver in resistance gene dissemination and contributor to the growing problem of antibiotic resistance globally.

*S. aureus* is a leading cause of many life-threatening infections and >90% of clinical isolates carry at least one plasmid (6). These plasmids carry a huge diversity of resistance genes and many encode for resistance to multiple classes of antibiotic (7). Conjugative plasmid transfer in *S. aureus* is initiated when a plasmid-encoded relaxase recognises, cleaves and attaches to the origin of transfer located on the plasmid. This nucleoprotein complex (known as the relaxasome) then directs the plasmid DNA to the mating pore, where it is transferred into the recipient cell through a type-IV secretion system (4). Mobilisable plasmids are plasmids that encode for all or part of the relaxasome but lack genes encoding for the mating pore. Therefore, they are dependent on a co-resident conjugative plasmid carrying compatible mating pore genes. Conjugative staphylococcal plasmids fall into three distinct families: the pSK41/pGO1 family, the pWBG749 family and the pWBG4 family (4). These families differ in their conjugation gene clusters. The pSK41/pGO1 family is by far the best studied and understood (8). pSK41 and pGO1 encode resistance to aminoglycosides, bleomycin and quaternary ammonium compounds. The two plasmids are essentially identical, but pGO1 contains a ∼8 kb co-integrated plasmid (encoding trimethoprim resistance) (9). The transfer region of pSK41/pGO1 contains most of the genes associated with conjugative plasmid transfer (8), while genes involved in replication, multimer resolution and plasmid partitioning are encoded in a second region, known as Region 1 (10). The conjugation frequency (*i.e.* the rate of transfer) of a given plasmid is controlled by two plasmid-specific factors: plasmid copy number (PCN), and the expression level of the conjugation machinery genes (*trs,* or transfer, genes) (11, 12). Logically, increased PCN and/or increased expression of transfer genes results in higher rates of plasmid transfer (13). The PCN of pSK41/pGO1 is controlled by the replication initiation protein, Rep (14). Because high rates of plasmid replication come with fitness costs for the host (15), expression of Rep is tightly controlled. The replication region of pSK41/pGO1, as well as encoding for Rep itself, encodes for two independent methods of Rep expression regulation. Firstly, expression of Rep is transcriptionally, but predominantly translationally, controlled by an antisense RNA (RNAI) that is complementary to the *rep* mRNA leader region (14, 16, 17). Secondly, the replication region also encodes for a small transcriptional repressor protein, Cop, that binds to the *rep* promoter and represses *rep* transcription (18). Expression of the transfer genes on pSK41/pGO1 is controlled by TrsN (known as ArtA in pSK41), a transcriptional repressor that binds to the promoters of *trsA, trsL* and three other promoters found within Region 1 (19, 20).

While conjugation frequency can be increased or decreased by plasmid mutations (11, 17), chromosomal mutations in the donor and environmental stimuli can also influence transmission rate (21, 22). In particular, stressors and the subsequently induced stress responses can promote increased conjugation frequency and mobilisation of MGEs more generally (23–26). Furthermore, conjugation is considered a stressful process for the donor and the induction of stress responses during mating is considered beneficial (27, 28). Induction of the SOS response has long been known to promote transmission of MGEs in both Gram-negatives and Gram-positives (25, 26). More specifically, SOS response induction by antibiotics and other pharmaceuticals leads to increased conjugation rates (22, 24, 29, 30) despite reducing the rate of plasmid replication (31). A second bacterial stress response – the stringent response – has also been implicated in elevated MGE transmission. Strugeon et al. observed that *Escherichia coli* growing in a biofilm had a higher rate of MGE excision than planktonic cells due to induction of the stringent response (23). Biofilms are often considered “hotspots” for horizonal gene transfer (32) and the stringent response is frequently implicated in biofilm formation (33). This stress response has also been postulated to play a role in MGE transfer in the plant symbiote *Mesorhizobium japonicum* (34). The stringent response is a conserved stress response mediated by the “alarmones” guanosine tetra- and pentaphosphate (collectively known as [p]ppGpp) (reviewed in (35)). In most bacteria, cellular levels of (p)ppGpp are mediated by a bifunctional enzyme, Rel, that both synthesises and hydrolyses (p)ppGpp *via* two distinct catalytic domains. Some bacteria, such as *S. aureus,* also possess accessory (p)ppGpp synthetases. In the classical description of the stringent response, limitation of one or more amino acid results in uncharged tRNAs entering the ribosome, resulting in stalling. This stalling is sensed by Rel, which responds by synthesising (p)ppGpp from ATP and GDP/GTP. The downstream effectors of (p)ppGpp vary greatly between species, with >50 known binding partners of (p)ppGpp (36). However, the overall effect of elevated (p)ppGpp is a downregulation of most metabolic processes and derepression of genes associated with amino acid synthesis and import. In *S. aureus*, the derepression effects of (p)ppGpp are mediated *via* CodY, a transcriptional repressor that requires GTP in order to bind to DNA (37). While the stringent response once thought to be binary, and either “on” or “off”, we now know that a basal level of (p)ppGpp is present in all cells and (p)ppGpp acts as a master regulator of almost all aspects of bacterial physiology and virulence (38).

Given the pervasive cellular effects of (p)ppGpp, it is not surprising that it has been found to play a role in various aspects of antibiotic susceptibility, resistance and killing (39–41). We have previously observed that Rel mutations identified in clinical isolates confer antibiotic tolerance in *S. aureus* by partially activating the stringent response (42, 43). Antibiotic tolerance describes the ability of a bacterial population to survive transient exposure to an otherwise lethal concentration of antibiotic without exhibiting resistance (44). It is emerging as an important contributor to persistent, relapsing and recurrent infections, as well as, worryingly, acting as a precursor to the development of endogenous resistance (40, 45). Given that plasmid conjugation is a major contributor to the spread of antibiotic resistance, and stress responses have been implicated in the transmission of MGEs, we set out to investigate whether clinical stringent response activation impacts plasmid transmission rates in *S. aureus*. Here we show that strains bearing clinical Rel mutations donate plasmids at significantly higher frequencies than wildtype strains. This effect applies to the transfer of diverse conjugative and mobilisable plasmids but, intriguingly, the molecular basis of this effect is neither increased PCN or expression of transfer genes. As Rel mutations are now being identified among further clinical isolates (46–48), this emerging relationship with plasmid transfer requires further investigation.

## RESULTS

### Rel mutation promotes pGO1 donation, but not receipt

We have previously introduced two clinical Rel mutations – F128Y and L152F – into the *S. aureus* model strain Newman and shown that they partially activate the stringent response by shifting the activity of Rel in favour of (p)ppGpp synthesis (42, 43). Combined evidence from growth curves, (p)ppGpp quantitation and enzyme assays suggests that the F128Y mutation induces a greater induction of the stringent response than L152F. In order to investigate the impact of these mutations on conjugal plasmid transfer, we began by measuring the conjugation frequency of pGO1 from SH1000 into novobiocin-resistant derivatives of wildtype Newman, Newman F128Y and a complemented strain (where the Rel F128Y mutation has been reversed). Because *S. aureus* conjugation requires cell-to-cell contact and growth in a biofilm (49), strains were combined and mated on nylon filters for four hours prior to plating out for donor, recipient and transconjugant quantification. There was no significant difference in the frequency of plasmid uptake by Newman, F128Y or the complemented strain (Figure 1A). However, when the matings were reversed and the Newman strains acted as pGO1 donors, the conjugation frequency for the F128Y mutant was ∼10-fold higher than that of the wildtype and complemented strains (Figure 1B). A similar, but smaller, effect was seen when the L152F mutant was used as the donor (Figure 1C). We confirmed that this effect was not due to a difference in donor-to-recipient ratio between strains; only data where the ratio fell between 0.5:1 and 1.5:1 were analysed, and collated data demonstrate that there is no correlation between conjugation frequency and donor-to-recipient ratio within this range (Figure S1). Therefore, stringent response activation in our Rel mutants promotes conjugal plasmid donation, but not receipt.

**Figure 1.**
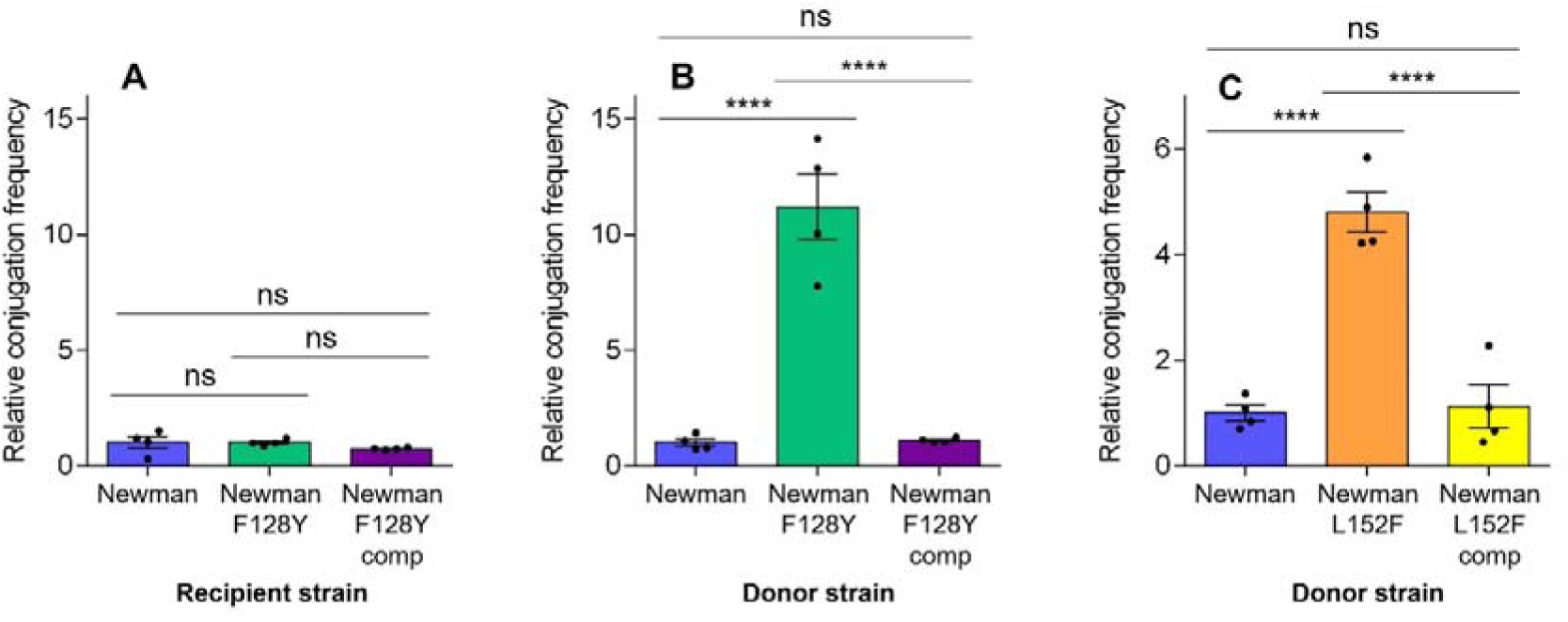
Clinical Rel mutations increase the frequency of pGO1 donation, but not receipt. (A) Filter matings were performed with SH1000 (pGO1) as donor and the novobiocin-resistant recipients indicated. (B) and (C) Filter matings were performed with different novobiocin-resistant donors (as indicated) carrying pGO1 and SH1000-NR as the recipient. All conjugation frequencies were calculated as transconjugants per donor and are expressed relative to the Newman mean. Data shown are the mean of four biological replicates; errors bars represent the SEM. Asterisks indicate statistically significant differences between means as determined by a one-way ANOVA with Tukey’s multiple comparisons test (**** = *P* ≤0.0001; ns = not significant).

### (p)ppGpp modulation affects the frequency of donation of diverse plasmids

In addition to mutation of Rel, the stringent response can be activated in bacteria by exposure to mupirocin, an isoleucyl-tRNA synthetase inhibitor. To corroborate our results with the Rel mutants, we exposed wildtype Newman carrying pGO1 to a subinhibitory concentration of mupirocin overnight prior to mating. Exposure to mupirocin induced a ∼4-fold increase in conjugation frequency compared with the unexposed control (Figure 2A). This effect of mupirocin, and the effect of the Rel F128Y mutation, on conjugation frequency is not strain-specific, as we observed the same trend with USA300 LAC (Figure S2). In *S. aureus*, while (p)ppGpp is primarily synthesised by Rel, there are two accessory (p)ppGpp synthetases that can contribute (RelP and RelQ). To test whether decreasing the cellular (p)ppGpp concentration would reduce plasmid donation, we generated a Δ*rel*_syn_ mutant of Newman that retains (p)ppGpp hydrolysis activity but lacks synthetic activity (Table S1) (50). In this Δ*rel*_syn_ background, we also deleted *relP* and *relQ* to generate a (p)ppGpp-null strain (51). Deletion of Rel synthetase activity led to a small (∼2-fold) but significant decrease in conjugation frequency compared with wildtype Newman (Figure 2B). When the (p)ppGpp-null mutant was used as the donor, the conjugation frequency was decreased slightly further. However, plasmid transmission still occurred, indicating that while elevated (p)ppGpp can promote conjugal plasmid transfer, it is not essential for conjugation.

**Figure 2.**
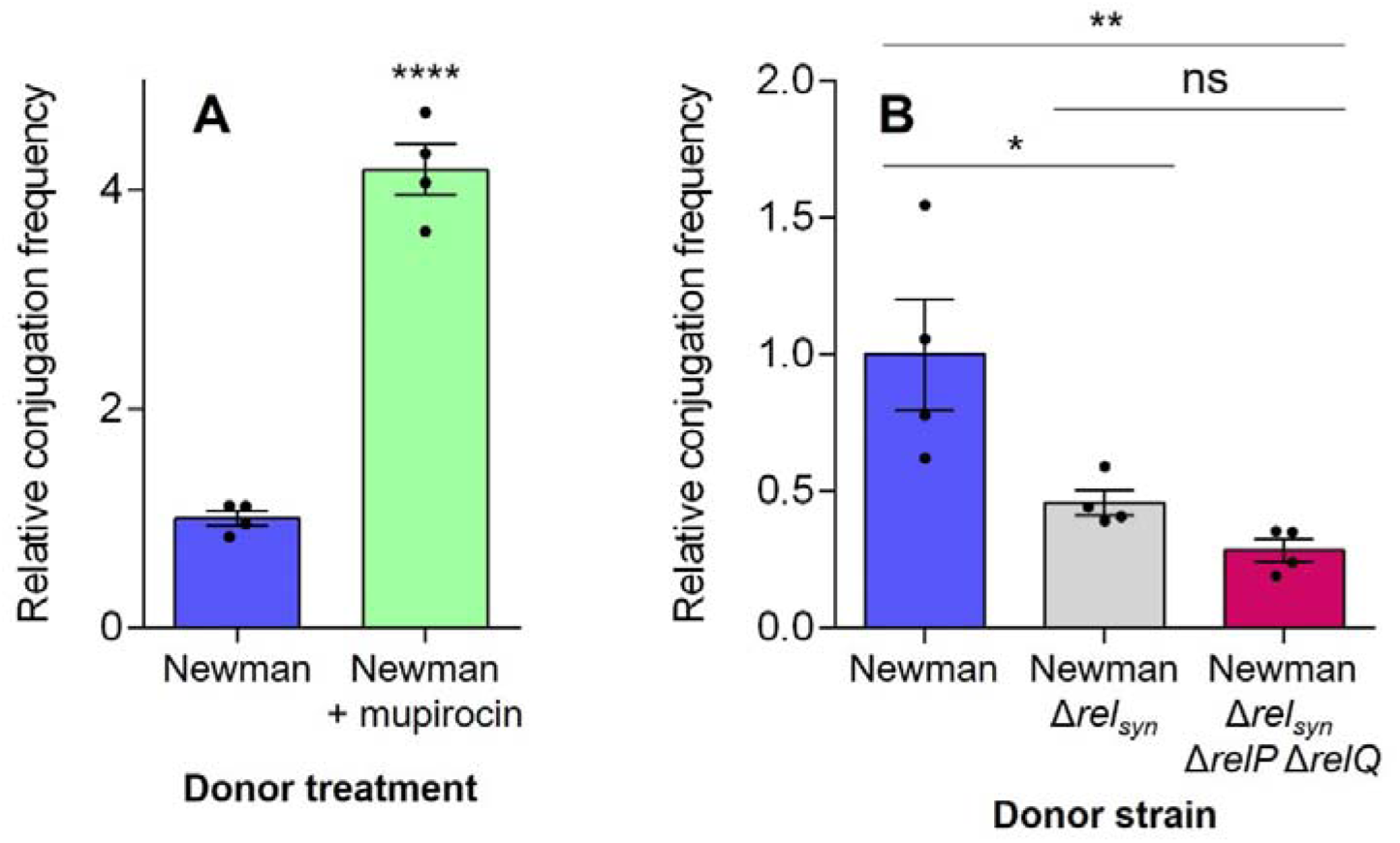
Modulation of (p)ppGpp level affects the frequency of pGO1 donation. Filter matings were performed with the novobiocin-resistant donor indicated and SH1000-NR as the recipient. (A) Conjugation frequency for transfer of pGO1 from Newman with and without chemical induction of the stringent response with mupirocin (donor was exposed to a subinhibitory concentration of mupirocin prior to mating). (B) Conjugation frequency for transfer of pGO1 from wildtype Newman and two stringent response-defective mutants. All conjugation frequencies were calculated as transconjugants per donor and are expressed relative to the Newman mean. Data shown are the mean of four biological replicates; errors bars represent the SEM. Asterisks indicate statistically significant differences between means as determined by a two-tailed *t*-test in panel A and a one-way ANOVA with Tukey’s multiple comparisons test in panel B (* = *P* ≤0.05; ** = *P* ≤0.01; **** = *P* ≤0.0001; ns = not significant).

Next, we decided to test whether the effect of stringent response activation on pGO1 conjugation frequency would also apply to other plasmids. pWBG707 and pWBG749e are representative conjugative resistance plasmids from the pWBG4 and pWBG749 families, respectively. They differ from pGO1 in their size (Table S1), resistances and conjugation genes (4). Conjugation of pWBG707 and pWBG749e from the F128Y mutant into SH1000 occurred at significantly higher frequencies than when Newman wildtype or the complemented strain were used as donors (although the effect was not as pronounced as with pGO1; Figure 3A and B). The F128Y mutant also mobilised pC221 when it was co-resident in cells with pWBG749e at a higher frequency than the wildtype and complement (Figure 3C). Therefore, the positive effect of stringent response activation on plasmid donation appears to be universal and not specific to the pSK41/pGO1 plasmid family.

**Figure 3.**
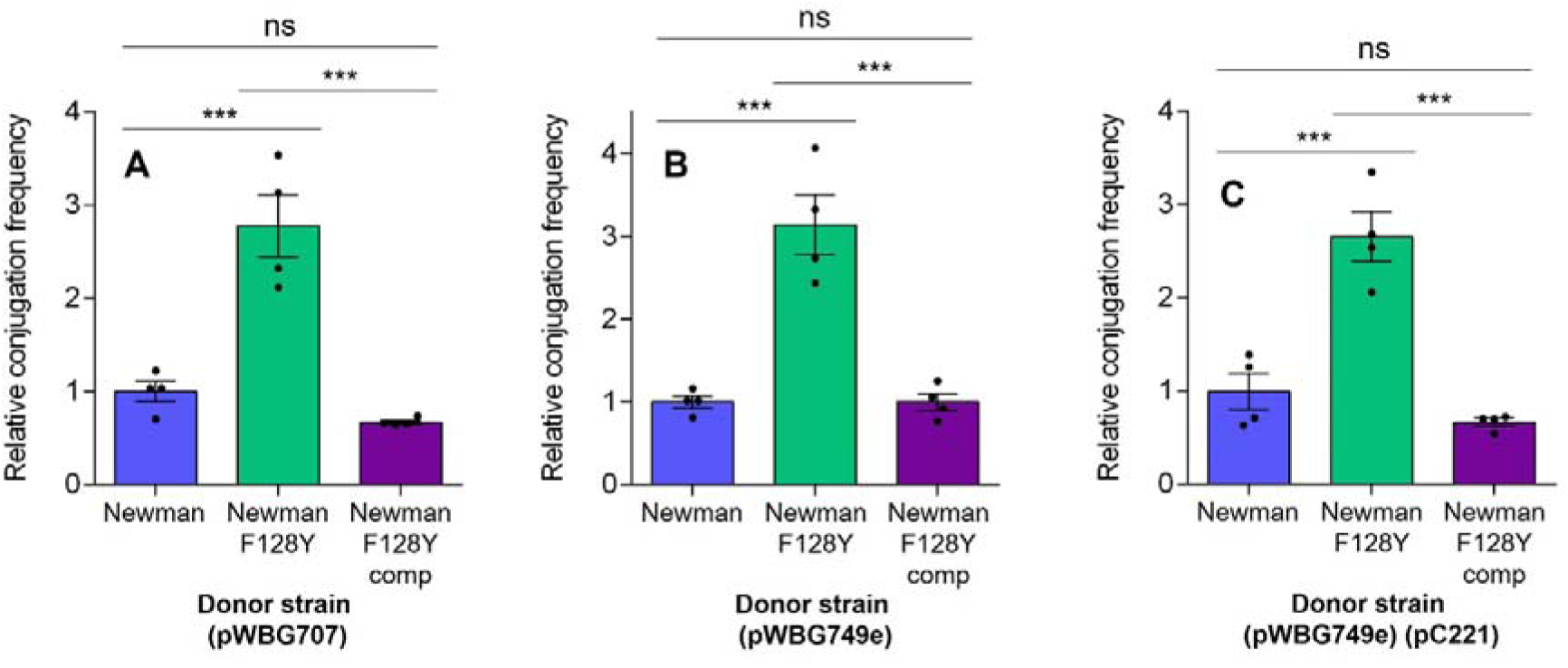
Clinical Rel mutation increases the donation frequency of diverse staphylococcal conjugative and mobilisable plasmids. Filter matings were performed with different novobiocin-resistant donors (as indicated) carrying different staphylococcal plasmids, and SH1000-NR as the recipient. (A) Conjugation frequency for transfer of pWBG707 from different Newman strains. (B) Conjugation frequency for transfer of pWBG749e from different Newman strains. (C) Mobilisation frequency for transfer of pC221 by pWBG749e from different Newman strains. All conjugation/mobilisation frequencies were calculated as transconjugants per donor and are expressed relative to the Newman mean. Data shown are the mean of four biological replicates; errors bars represent the SEM. Asterisks indicate statistically significant differences between means as determined by a one-way ANOVA with Tukey’s multiple comparisons test (*** = *P* ≤0.001; ns = not significant).

### (p)ppGpp does not promote plasmid donation *via* CodY derepression, induction of the SOS response or increased plasmid copy number

Next, we were interested to investigate the molecular mechanism behind the (p)ppGpp and plasmid conjugation relationship. As mentioned previously, part of stringent response activation in *S. aureus* is release of the transcriptional repressor CodY from transcripts. To test whether CodY derepression may play a role in increased plasmid transfer by stringent response-activated donors, we compared the conjugation frequency of pGO1 between wildtype Newman and a *codY* deletion mutant as donors (Figure 4A). The Δ*codY* strain exhibited a slightly higher conjugation frequency than the wildtype, but the difference was not statistically significant and not on the same scale as the F128Y mutant or mupirocin treatment. We also tested the potential contribution of the SOS response. The stringent and SOS responses are intrinsically linked (52) and a previous study found that increased MGE transmission upon stringent response induction was due to downstream activation of the SOS response (23). The SOS response is mediated by the repressor LexA, which must undergo self-cleavage for the SOS response to be induced. Therefore, mutation of the catalytic serine (S130) in LexA prevents SOS induction (53). We generated a LexA S130A mutant of Newman and exposed it to mupirocin, to test whether a functional SOS response is required for (p)ppGpp-stimulated plasmid donation (Figure 4B). The LexA mutant exhibited a large and significant increase in conjugation frequency following mupirocin exposure, indicating that the SOS response is not necessary for the relationship between (p)ppGpp and plasmid transfer.

**Figure 4.**
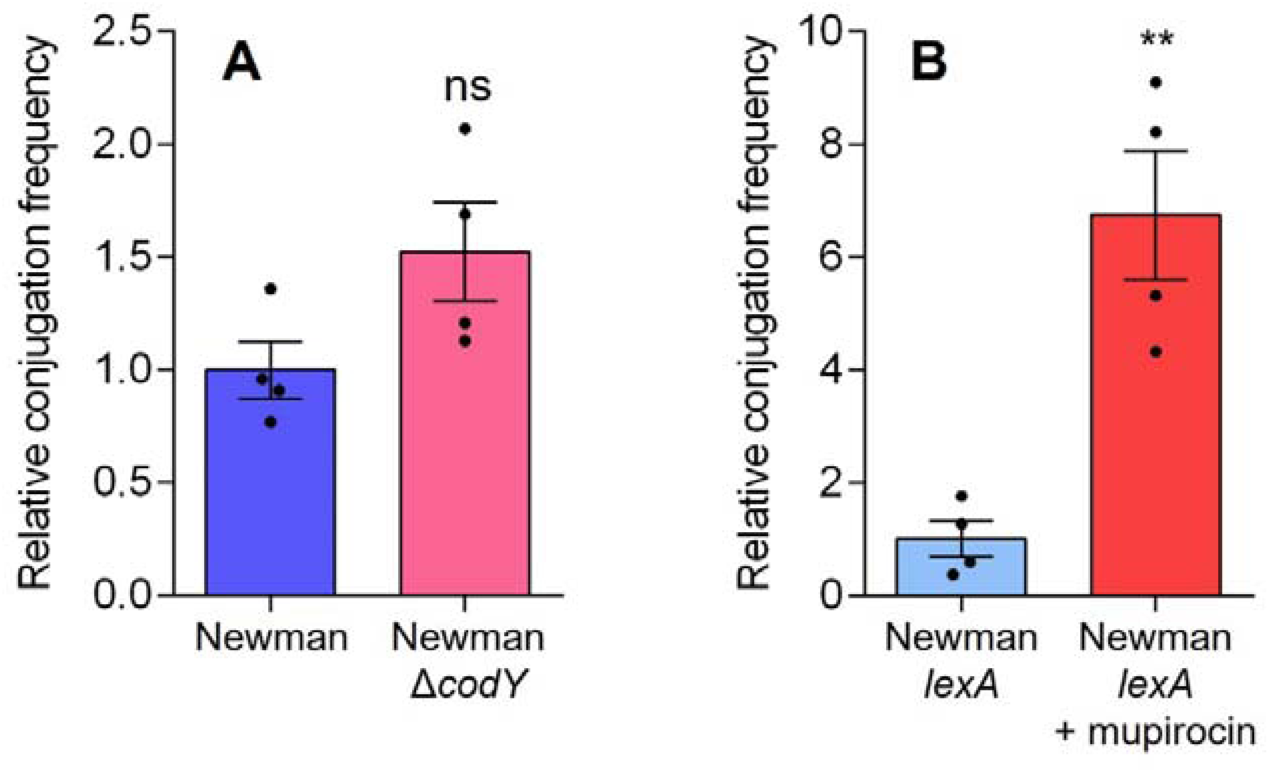
(p)ppGpp does not promote donation of pGO1 *via* CodY derepression or induction of the SOS response. Filter matings were performed with the novobioin-resistant donor indicated and SH1000-NR as the recipient. (A) Conjugation frequency for transfer of pGO1 from wildtype Newman and a Δ*codY* mutant. (B) Conjugation frequency for transfer of pGO1 from a *lexA* mutant with and without chemical induction of the stringent response with mupirocin (donor was exposed to a subinhibitory concentration of mupirocin prior to mating). All conjugation frequencies were calculated as transconjugants per donor and are expressed relative to the wildtype Newman or no mupirocin mean. Data shown are the mean of four biological replicates; errors bars represent the SEM. Asterisks indicate statistically significant differences between means as determined by a two-tailed *t*-test (** = *P* ≤0.01; ns = not significant).

Increased conjugation frequency can occur due to elevated PCN, and PCN can be phenotypically reflected in the level of resistance of a strain (11). Therefore, we compared the pGO1-encoded resistance of wildtype Newman, the F128Y mutant and the complemented strain. Interestingly, the wildtype and complemented strains had minimum inhibitory concentrations (MICs) for gentamicin of 50 and 75 µg/mL, respectively, while the F128Y mutant had a reproducible MIC of 200 µg/mL. The higher gentamicin resistance phenotype of F128Y compared with the wildtype was also confirmed by population analysis profile determination (Figure S3). However, when the MIC for trimethoprim was determined, the F128Y mutant had an MIC of 1,400 µg/mL while the wildtype and complemented strains had MICs of ≥1,500 µg/mL.

To further explore whether PCN is elevated in the F128Y mutant, two reporter constructs were employed. Firstly, a PCN reporter construct for pGO1/pSK41 has previously been reported (17). pSK5487 is a mini replicon encoding for chloramphenicol acetyltransferase (CAT) and whose replication in *S. aureus* is dependent on the pGO1/pSK41 replication region. We introduced pSK5487 into wildtype Newman, F128Y and the complemented strain and quantified CAT activity in cell lysates. As elevated (p)ppGpp is associated with a downregulation of protein translation, we expressed the CAT activity as per ng of total protein. No significant difference in CAT activity was observed between the three strains (Figure 5A). As the PCN of pGO1 is regulated by Rep and the antisense RNAI, we also constructed a reporter construct in which the expression of LacZ is under the control of the complete pGO1 *rep* promoter region and introduced this into our three strains. Again, we saw no significant difference in reporter enzyme activity between strains (Figure 5B). Finally, to provide a measure of absolute PCN that would not be potentially confounded by the metabolic downregulation mediated by elevated (p)ppGpp, we performed droplet digital PCR on plasmid DNA extracted from our three strains (Figure 5C). The PCN of pGO1 in wildtype Newman was determined to be approximately 5, which is consistent with other PCN estimates for similarly large plasmids (54). The PCN of pGO1 in F128Y was not statistically significantly different from that of wildtype Newman or the complemented strain. Therefore, we can conclude from this body of evidence that elevated PCN is not the molecular basis of increased conjugation frequency in our stringent response-activated strains.

**Figure 5.**
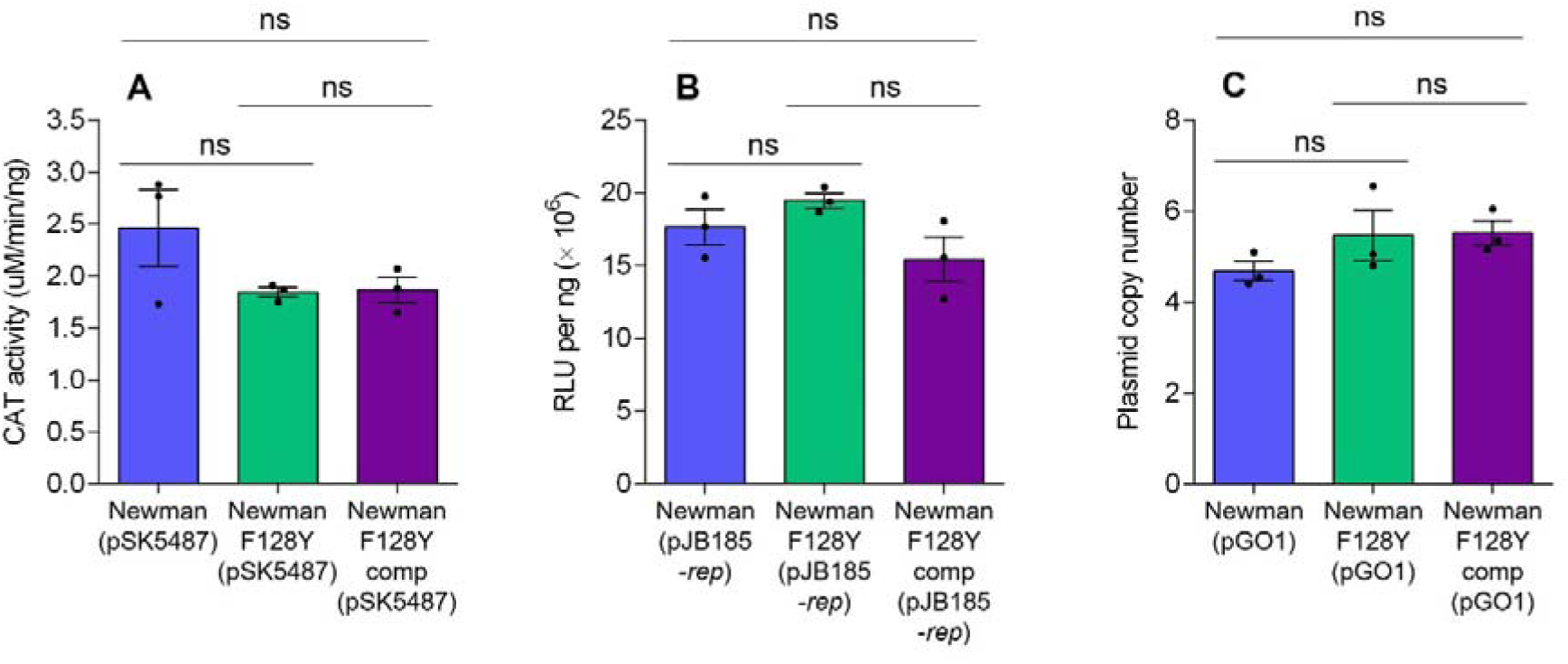
Clinical Rel mutation does not increase pGO1 copy number. (A) Chloramphenicol acetyltransferase (CAT) activity in different strains carrying pSK5487, a plasmid copy number reporter construct bearing the pGO1 replication region. (B) LacZ activity in different strains carrying pJB185-*rep*, a reporter construct in which expression of *lacZ* is under the control of the pGO1 *rep* promoter region. In both panels, data were corrected for total protein content and are derived from three independent replicate cultures per strain (and two technical replicates). (C) Absolute plasmid copy number of pGO1 for each strain as determined by droplet digital PCR. Data shown are the mean of three independent biological replicates and two technical replicates. In all panels, errors bars represent the SEM. statistically significant differences between means were assessed by a one-way ANOVA with Tukey’s multiple comparisons test (ns = not significant).

### Transcriptomic profile of F128Y (pGO1) fails to reveal mechanism of increased conjugal plasmid transfer

Our data indicate that PCN is not altered in our F128Y mutant. However, conjugation frequency can also be elevated when expression of the plasmid transfer genes is increased (11, 19). The repressor TrsN binds to several transfer gene promoters, including that of *trsA* (19). Therefore, we constructed a *trsA* expression reporter, where the *trsA* promoter drives expression of LacZ. When this construct was introduced into Newman wildtype and the F128Y mutant with and without pGO1, the repressive effect of TrsN expressed from pGO1 on the expression of *lacZ* was clear (Figure 6). However, there was no significant difference in LacZ activity between the wildtype and mutant.

**Figure 6.**
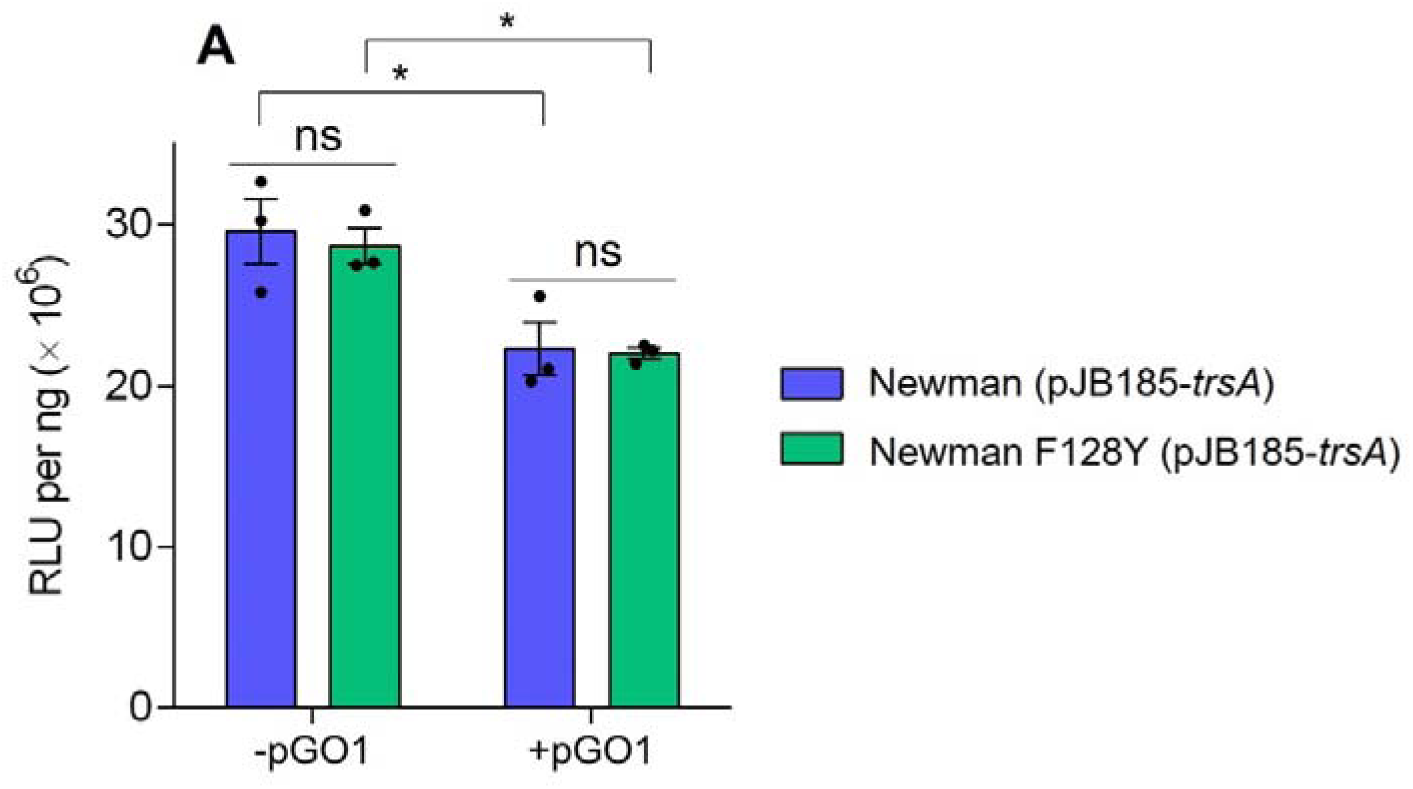
Clinical Rel mutation does not affect the expression of plasmid-encoded genes, including conjugation machinery genes. (A) LacZ activity in different strains carrying pJB185-*trsA (*a reporter construct in which expression of *lacZ* is under the control of the pGO1 *trsA* promoter) and in the presence/absence of pGO1. Data were corrected for total protein content and are derived from three independent replicate cultures per strain (and two technical replicates). Errors bars represent the SEM. Asterisks indicate statistically significant differences between means as determined by a one-way ANOVA with Tukey’s multiple comparisons test (* = *P* ≤0.05; ns = not significant).

To directly measure the expression of all plasmid-encoded genes, we performed RNA sequencing on wildtype Newman and the F128Y mutant each bearing pGO1. Of the 57 transcripts originating from pGO1, only one was differentially expressed (log2 fold change >1 [equal to >2-fold change] and adjusted *p*-value <0.05), with 3.5-fold greater expression in wildtype Newman: MobV family relaxase (locus tag PGO1_p19). Plasmid-encoded relaxases are involved in cleaving plasmid DNA at the *oriT* and initiating conjugative transfer; however, PGO1_p19 is not the canonical and well-studied pGO1/pSK41 relaxase, NES, that is encoded next to *oriT* and considered to be part of the core pGO1 conjugative replicon (55, 56). Furthermore, downregulation of a relaxase in the F128Y mutant does not intuitively align with *increased* conjugation frequency. If the threshold for differential expression is lowered to a 1.5-fold change, expression of NES was also higher in the wildtype than the mutant (as well as two neighbouring genes encoding for small hypothetical proteins on the opposite strand). With this lower fold-change threshold, two genes were more highly expressed in the mutant than the wildtype: a putative resolvase, known as Res (57), and a 55 amino acid hypothetical protein. Plasmid-encoded resolvases act to resolve plasmid dimers into monomers during partitioning and contribute to plasmid maintenance (58). We could find no precedent for a relationship between resolvase activity and conjugation frequency in the literature. However, overlap between tolerance mutations and increased plasmid maintenance has been reported in *Escherichia coli* (59). Manual inspection of all reads that mapped to pGO1 – including antisense and intergenic regions – revealed no other differences between the two datasets.

Based on a >2-fold change and adjusted *p*-value <0.05, 96 chromosomal genes were upregulated in the F128Y mutant and 163 genes were downregulated. The most striking differential expression was the upregulation of capsule biosynthesis genes in the F128Y mutant (2.5-3.5-fold change, adjusted *p*-value <0.05). This same property was observed in the original clinical isolate bearing the F128Y mutation *via* microarray analysis (60) and these genes are part of the CodY regulon (61). Capsule type and volume have been shown to impact conjugation efficiency in Gram-negative bacteria (where capsule volume is negatively correlated with conjugation efficiency) (62). However, we observed elevated conjugation frequency in a *rel* F128Y mutant of USA300 LAC (Figure S2), which lacks a capsule (63). Therefore, the upregulation of capsule biosynthesis genes in the F128Y mutant is unlikely to be the cause of elevated conjugation frequency. No other significant changes in gene expression in the mutant (either up or down) were noted that have previously been linked to conjugation frequency (*e.g.* biofilm formation, cell membrane composition and cell permeability (22, 49, 64)).

## DISCUSSION

The stringent response is a master regulator of many bacterial physiological processes – from growth, metabolism and sporulation to biofilm formation, virulence expression and antibiotic tolerance (38, 39). The results presented here indicate that plasmid conjugation should be added to this list. Using two mutant strains that carry clinical Rel mutations, as well as chemical and further genetic activation of the stringent response, we have shown that elevated (p)ppGpp levels lead to higher rates of plasmid donation. This effect applies to plasmids from the three different families of staphylococcal plasmid, in addition to a mobilisable plasmid. Our comprehensive experimental investigations into plasmid-mediated factors (*i.e.* PCN and conjugation machinery expression) revealed no differences between wildtype and mutant. Transcriptomic analysis of pGO1 also revealed only small changes in a limited number of genes, none of which could be speculated to cause the difference in conjugation frequency. Therefore, we must consider host factors. An obvious difference between our Rel mutants and the wildtype is lag time and/or growth rate (42). However, higher rates of plasmid transfer are associated with *faster* growing bacteria (65, 66), so the slow growth of our Rel mutants does not explain our findings. Many host factors have been linked to differences in conjugation frequency, including cell permeability, cell envelope changes and a range of metabolism-associated genes (21, 22, 64, 67). While we were unable to match any of these reported associations with our host transcriptomic analysis, the impact of the stringent response on gene expression is pervasive; at its peak, (p)ppGpp can regulate up to a quarter of the genome (68). (p)ppGpp also has significant post-translational effects, binding to a range of proteins and enzymes, modifying their activity (36). Given that we did not see any difference in conjugation frequency when the recipient had an activated stringent response, the effect is clearly occurring within the donor. Overall, further work is required to determine the molecular basis of increased plasmid donation associated with stringent response activation.

The implications of this new finding are significant. Mutations in Rel and accessory (p)ppGpp synthetases are increasingly being identified among clinical isolates (46–48, 60), where they already confer antibiotic tolerance and impact the expression of resistance (42, 43). Our findings suggest that these mutations can also promote the dissemination of resistance plasmids, contributing to the wider problem of resistance. Due to the role of the stringent response in bacterial growth, stress survival and virulence, it is already under investigation as a target for novel antibiotics (69). The emerging relationship between the stringent response and plasmid dissemination reported here adds further impetus to this search, with the potential to prevent plasmid dissemination, reverse antibiotic tolerance and reduce virulence with one therapeutic.

## METHODS

### Reagents and antibiotics

X-gal (5-bromo-4-chloro-3-indolyl-β-D-galactopyranoside) and chloramphenicol were purchased from Bio Basic Inc. (Markham, ON, Canada). All other antibiotics and reagents, unless otherwise stated, were purchased from MilliporeSigma/Merck.

### Bacterial strains, plasmids and growth conditions

All bacterial strains and plasmids used in this study are listed in Table S1. *S. aureus s*trains were routinely cultured in tryptic soy broth (TSB) and on tryptic soy agar (TSA) at 37 °C, while *E. coli* strains were cultured in/on Lysogeny Broth (LB) or agar. The Newman *rel* mutants F128Y and L152F and their complemented counterparts (F128Y comp and L152F comp) have been reported previously (42), as has the *rel* F128Y mutant of USA300 LAC (43). Strains were made novobiocin-resistant (NOV^R^) as previously described (43). SH1000 was made novobiocin- and rifampicin-resistant by plating on TSA containing 1 μg/mL novobiocin and 0.5 μg/mL rifampicin, while Newman F128Y comp was made resistant to fusidic acid by plating on TSA containing 1 μg/mL fusidic acid.

### Mutant generation *via a*llelic exchange

To delete residues 308-310 from Rel (GenBank accession number NWMN_RS08620) and inactivate the synthetase domain, the two flanking halves of the gene were amplified from Newman genomic DNA and joined together by overlap extension PCR. For the *relP* and *relQ* knockouts, left and right flanks of each gene (and surrounding genes when needed to give fragments >500 bp) were amplified and joined together, excluding an internal fragment that contains conserved domains responsible for synthetic activity (51). For *codY*, a complete deletion of the gene (except the start and stop codons) was generated by amplifying the up- and downstream genes (and intergenic regions) and joining together. The *lexA* S130A insert was generated by amplifying at least 500 bp up- and downstream of the mutation site with mutagenic primers and joining together. All inserts were cloned into pIMAY-Z (70) between the EcoRI and NotI sites *via* In-Fusion cloning (TaKaRa Bio), transformed into *Escherichia coli* Stellar, and plated on LB agar containing 10 µg/mL chloramphenicol and 50 µg/mL X-gal at 37 °C. Confirmed pIMAY-Z constructs were transformed into *E. coli* IM08B (70), extracted, electroporated into competent *S. aureus* cells, and integrated/excised as previously described (42). Mutants were confirmed initially by PCR and bidirectional sequencing, and latterly by long-read whole genome sequencing (Novogene). All primers are listed in Table 2. For construction of Δ*rel_syn_*Δ*relP* Δ*relQ*, the *rel*_syn_ mutation was introduced first, followed by the *relP* deletion and then the *relQ* deletion.

### Introduction of plasmids into donor strains *via* conjugation

Conjugative plasmids were introduced into the different NOV^R^ donor strains *via* conjugation with NOV-sensitive hosts. Unmarked SH1000 (pGO1) was used to conjugate pGO1, while WBG541 (71) was used as the source of pWBG749e (72). pWBG707 (73) was provided in a NOV^R^ strain, WBG4515 (74), so pWBG707 was first conjugated into Newman F128Y comp FUS^R^ before being transferred into the NOV^R^ donor strains. To introduce both pGO1 and pC221 (75) into NOV^R^ donor strains, pGO1 was first conjugated into RN4220 (pC221) and the transconjugant was then used to introduce both plasmids into the NOV^R^ strains simultaneously. These conjugations were performed essentially as described below, except matings were incubated for 24 hours and no quantitation of donor and recipient was performed. Transconjugants were restreaked on drug plates to ensure purity. Antibiotic concentrations used to select for plasmids and chromosomal resistance were: gentamicin (5 μg/mL); trimethoprim (10 μg/mL); erythromycin (5 μg/mL); chloramphenicol (10 μg/mL); novobiocin (1 μg/mL); rifampicin (5 μg/mL) and streptomycin (50 μg/mL).

### Conjugation/mobilisation frequency determinations

Conjugation and mobilisation frequencies were determined as described by Savage et al. (49) with some modifications. Donor and recipient strains were grown on selective agar at 37 °C overnight then resupended in antibiotic-free TSB to an OD_600nm_ of 1.0. Where chemical stringent response induction was required, mupirocin at 0.04 μg/mL was included in the agar plate. When acting as the donor, wildtype and complemented strains were combined with the recipient at a 1:2 ratio (250 μL wildtype/complemented and 500 μL SH1000-NR), while mutant strains were combined with SH1000-NR at a 1:1 ratio (500 μL each). This ensured a consistent donor-to-recipient ratio between strains. Mating mixtures were pushed through a syringe onto a 0.2 μM pore-size nylon filter using 13 mm Swinnex filter holders. Filters were placed bacteria side down on TSA and incubated at 37 °C for 4 hours. Following incubation, cells were resuspended in 1 mL TSB and serially diluted in PBS. Serial dilutions were plated on TSA containing the appropriate antibiotics for selection of donors, recipients or transconjugants, and incubated at 37 °C for 16-24 hours. Colonies were enumerated either manually or using an aCOLyte 3 automated colony counter (Synbiosis Ltd., Cambridge, UK). Four independent matings were performed per donor-recipient combination/condition. The donor-to-recipient ratio was calculated for each mating and any matings where the ratio fell below 0.5:1 or above 1.5:1 were excluded. Conjugation/mobilization frequencies were calculated as transconjugants per donor.

### Antibiotic susceptibility assays

Minimum inhibitory concentrations (MICs) were determined in triplicate in Mueller-Hinton broth (MHB) following the broth microdilution guidelines of the Clinical and Laboratory Standards Institute (CLSI) (76). Population analysis profiles with gentamicin were determined by growing the test strains overnight in TSB, diluting in phosphate-buffered saline, and plating on TSA containing increasing concentrations of gentamicin. Plates were incubated for 24 hours prior to colony counting. Three independent overnight cultures per strain were plated out.

### Reporter construct generation

The staphylococcal reporter vector pJB185(77) was amplified and linearised by PCR using primers listed in Table 2, and the methylated template DNA degraded with DpnI. The pGO1 *rep* promoter region (including RNAI) and the *trsA* promoter were amplified from a lysed colony of SH1000 (pGO1). Inserts were cloned into pJB185 *via* In-Fusion cloning, transformed into *E. coli* Stellar and plated on ampicillin. Following confirmation of insert by bidirectional sequencing, purified plasmids were transformed into *E. coli* IM08B, extracted and introduced into *S. aureus* strains either by electroporation or phage transduction with phi85 (78).

### Chloramphenicol acetyltransferase (CAT) reporter assay

The plasmid copy number reporter construct pSK5487 was introduced into Newman strains *via* phage transduction. For the reporter assay, the method of Kwong et al. (14) was followed with a few modifications. Briefly, samples of overnight cultures grown in TSB were pelleted, resuspended in lysis buffer containing 0.13 mg/ml lysostaphin and incubated at 37 °C for 30 mins prior to sonication. Insoluble material was then pelleted and the supernatants diluted 1/10 prior to use. For the CAT assay, 93 μL CAT assay mixture (100 mM Tris, pH 7.8, 0.1 mM acetyl-CoA, 1 mM 5,5′-dithiobis-(2-nitrobenzoic acid) and 5 μL diluted cell supernatant were combined in a 96-well plate and equilibrated to 37 °C for 2 mins in a Tecan Spark plate reader. The reaction was initiated with 2 μL 5 mM chloramphenicol, shaken for 5 seconds and the absorbance at 415 nm monitored. The initial rate of CAT activity was determined from the slope and converted to μM/min using the extinction coefficient of the product 5-thio-2-nitrobenzoic acid (14150 M/cm) and a pathlength of 0.28 cm. The total protein content of each cell supernatant was determined using a Bradford assay and this value used to normalise the rate of CAT activity. Three independent cultures were grown, lysed and assayed per strain.

### LacZ reporter assays

The *rep* and *trsA* reporter constructs pJB185-*rep* and pJB185-*trsA* were introduced into Newman strains *via* phage transduction. Cell extracts were prepared as described for the CAT report assay, except that cells from mid-exponential phase were pelleted and lysed. The Beta-Glo® assay system (Promega) was used to quantify LacZ expression in 50 μL diluted cell supernatants following the manufacturer’s instructions. Total protein content of each cell supernatant was determined as described above and used to normalise the luminescence values. Three independent cultures were grown, lysed and assayed per strain.

### Droplet digital PCR

Droplet digital PCR (ddPCR) was used for the absolute quantification of pGO1 PCN in Newman Wildtype, F128Y and F128Y comp. For these assays, triplicate 2 mL overnight cultures of each strain grown in TSB (supplemented with gentamicin) were harvested by centrifugation, cells washed twice with PBS and resuspended in 1 mL of PBS. The suspension was subjected to bead-beating using the standard B-matrix protocol and a FASTPREP-24 5G instrument (MP Biomedicals, USA). The homogenate was centrifuged for 5 minutes at 13,000 rpm, and the supernatant containing total DNA retained. One microliter of a 10⁻□ dilution in sterile water was used as the template for ddPCR. Each ddPCR reaction (25 µL total volume) contained nuclease-free water, 2× EvaGreen ddPCR Supermix (Bio-Rad) and primers at a final concentration of 0.1 µM. Primer sequences for amplification of *femA* (chromosomal; NWMN_RS07245) (79) and *aphD* (pGO1 plasmid; PGO1_RS00250) fragments are provided in Table S2. No-template controls (NTCs) were included to monitor contamination and primer–dimer formation. Reactions were dispensed in duplicate into a 96-well semi-skirted PCR plate. Plates were heat-sealed with pierceable foil, vortexed briefly, centrifuged for 1 minute, and loaded into an AutoDG instrument (Bio-Rad) for droplet generation using EvaGreen AutoDG oil. The resulting droplet plate was heat-sealed and transferred to a C1000 Touch Thermal Cycler (Bio-Rad). PCR amplification was performed according to the manufacturer’s instructions under the following cycling conditions: 95 °C for 2 min (1 cycle); 30 cycles of 94 °C for 30 s (ramp rate 2 °C/min) and 50 °C for 30 s (ramp rate 2 °C/min), 72°C for 2 min; followed by 70 °C for 10 min, and a final hold at 8 °C until droplet reading. After amplification, the plate was loaded onto a QX200 Droplet Reader (Bio-Rad) and analyzed using QuantaSoft software version 1.7.4 (Bio-Rad). Each droplet functions as an independent PCR reaction, and fluorescence detection distinguishes positive from negative droplets. For each well, the concentration of target molecules was calculated based on the fraction of positive droplets assuming a Poisson distribution. The associated error for each well was reported as the 95% confidence interval derived from the Poisson distribution, reflecting the statistical uncertainty of target molecule partitioning into droplets. PCN was calculated as the ratio of (copies/µL plasmid gene) to (copies/µL chromosomal gene). PCN values represent the mean of two technical replicates derived from three independently prepared DNA samples per strain.

### RNA sequencing

Cells for RNA extraction were grown exactly as described for conjugation frequency determinations. Following the 4-hour incubation, filters were removed from plates, the cells resuspended in 1 mL TSB and combined with 2 volumes of RNAprotect (Qiagen). From here, total RNA samples were prepared, quantified and sequenced by Novogene as described previously (80). RNA was extracted from four independent filter matings per donor. Gene expression was quantified using Salmon v1.10.2 (81) by pseudoaligning reads to CDS from NC_009641.1 (82) and NC_012547 (9). The resulting counts were modelled as a function of strain using DESeq2 1.40.2 (83). Differential gene expression was determined by a threshold of absolute fold change greater than 2 and an FDR-adjusted *p* value less than 0.05. Manual inspection of strand-specific aligned reads was performed in Artemis (84). Raw sequencing data has been deposited under BioProject accession number PRJNA1349606.

## ACKNOWLEDGEMENTS

We thank Josh Ramsay (Curtin University) for providing plasmids and useful discussions. We also thank Alex O’Neill (University of Leeds), Stephen Kwong (Western Sydney University) and Jeffrey Bose (University of Kansas) for providing plasmids. This work was supported by a project grant from the Canadian Institutes of Health Research (CIHR) awarded to ABB and JKH PJT173349), and a research grant from the Royal Society awarded to JKH (RG\R2\232237). ATD was supported by a CIHR Doctoral Foreign Study Award. Acquisition of the QX200 AutoDG (Automated Droplet Generator) ddPCR system was supported by the Polish Ministry of Science and Higher Education under the Capital Equipment Grant No. 6830/IA/SP/2018 awarded to A-KK. The authors acknowledge Research Computing at the James Hutton Institute for providing computational resources and technical support for the “UK’s Crop Diversity Bioinformatics HPC” (BBSRC grants BB/S019669/1 and BB/X019683/1), use of which has contributed to the results reported within this paper.

