## Supplemental Table 1, Supplemental Table 2, Supplemental Figure 1, Supplemental Figure 2, Supplemental Figure 3 for "Clinical stringent response activation promotes conjugal transfer of staphylococcal resistance plasmids"

**Table S1. List of bacterial strains and plasmids used in this study**

| **Bacterial strain/plasmid** | **Description** | **Source/Reference** |
| --- | --- | --- |
| **Bacterial strains** | | |
| Newman NOV^R^ | Novobiocin-resistant version of wildtype Newman bearing *gyrB* R144I mutation | Deventer et al. (43) |
| Newman *rel* F128Y NOV^R^ | Novobiocin-resistant version of Newman *rel* mutant bearing *gyrB* R144I mutation | Deventer et al. (43) |
| Newman *rel* F128Y comp NOV^R^ | Novobiocin-resistant version of complemented Newman *rel* mutant bearing *gyrB* R144I mutation | This study |
| Newman *rel* F128Y comp FUS^R^ | Fusidic acid-resistant version of complement Newman *rel* mutant | This study |
| Newman *rel* L152F NOV^R^ | Novobiocin-resistant version of Newman *rel* mutant bearing *gyrB* R144I mutation | Deventer et al. (43) |
| Newman *rel* L152F comp NOV^R^ | Novobiocin-resistant version of complemented Newman *rel* mutant bearing *gyrB* R144I mutation | This study |
| USA300 LAC NOV^R^ | Novobiocin-resistant version of wildtype USA300 LAC bearing *gyrB* R144I mutation | This study |
| USA300 LAC *rel* F128Y NOV^R^ | Novobiocin-resistant version of USA300 LAC *rel* mutant bearing *gyrB* R144I mutation | This study |
| Newman Δ*rel_syn_* | Newman mutant bearing Δ924-930 deletion in Rel synthetase domain | This study |
| Newman Δ*rel_syn_* NOV^R^ | Novobiocin-resistant version of Newman Δ*rel_syn_* bearing *gyrB* R144I mutation | This study |
| Newman Δ*rel_syn_* Δ*relP* Δ*relQ* | Newman mutant bearing Δ924-930 deletion in Rel synthetase domain and deletions of *relP* and *relQ* | This study |
| Newman Δ*rel_syn_* Δ*relP* Δ*relQ* NOV^R^ | Novobiocin-resistant version of Newman Δ*rel_syn_* Δ*relP* Δ*relQ* bearing *gyrB* R144I mutation | This study |
| Newman Δ*codY* | Newman mutant bearing *codY* deletion | This study |
| Newman Δ*codY* NOV^R^ | Novobiocin-resistant version of Newman Δ*codY* bearing *gyrB* R144I mutation | This study |
| Newman *lexA* S130A | Newman mutant bearing S130A mutation in *lexA* | This study |
| Newman *lexA* S130A NOV^R^ | Novobiocin-resistant version of Newman *lexA* S130A bearing *gyrB* R144I mutation | This study |
| RN4220 | Restriction-deficit laboratory strain | Fairweather et al. (85) |
| SH1000 | Standard laboratory strain | Horsburgh et al. (86) |
| SH1000-NR | Novobiocin- and rifampicin-resistant version of SH1000 | This study |
| WBG541 | Fusidic acid- and rifampicin-resistant laboratory strain | Udo et al. (71) |
| WBG4515 | Streptomycin- and novobiocin-resistant laboratory strain | Townsend et al. (74) |
| **Plasmids** | | |
| pGO1 | Prototypical conjugative staphylococcal plasmid. Encodes for gentamicin and trimethoprim resistance. Member of pGO1/pSK41 plasmid family. 54 kb. | Dr Alex O’Neill, University of Leeds  Caryl & O’Neill (9) |
| pWBG707 | Conjugative staphylococcal plasmid. Encodes for trimethoprim resistance. Member of the pWBG4 plasmid family. 38 kb. | Dr Josh Ramsay, Curtin University  Udo et al. (73) |
| pWBG749e | Erythromycin-resistant derivative of conjugative staphylococcal plasmid pWBG749 carrying Tn551. Member of the pWBG749 plasmid family. 43 kb. | Dr Josh Ramsay, Curtin University  O’Brien et al. (72) |
| pC221 | Mobilisable staphylococcal plasmid. Encodes for chloramphenicol resistance. | Dr Alex O’Neill, University of Leeds  Projan et al. (75) |
| pSK5487 | Plasmid copy number *cat* reporter construct carrying pGO1/pSK41 replication region. | Dr Stephen Kwong, Western Sydney University  Kwong et al. (14) |
| pJB185 | *S. aureus lacZ* reporter construct. | Dr Jeffrey Bose, University of Kansas  Krute et al. (77) |
| pJB185-*rep* | pGO1 *rep* promoter-*lacZ* fusion construct | This study |
| pJB185-*trsA* | pGO1 *trsA* promoter region-*lacZ* fusion construct | This study |

**Table S2. Primers used in this study**

| **Primer name** | **Use** | **Primer sequence (5’→3’)** |
| --- | --- | --- |
| *rel* upstream fwd | Amplification of *rel* gene half | ATAAGCTTGATATCGATGAAC  AACGAATATCCATATAGTGC |
| *rel* downstream rev | Amplification of *rel* gene half | ACCGCGGTGGCGGCCCTAG  TTCCAAACTCTTGTTACTGTATAAAC |
| *rel*-syn upstream rev | Amplification of *rel* gene half lacking nucleotides 924-930 | TACTGTAGTATGCAACAAATT  TTGTTTAGGCATTGCAATATAATC |
| *rel*-syn downstream fwd | Amplification of *rel* gene half lacking nucleotides 924-930 | CCTAAACAAAATTTGTTGCAT  ACTACAGTAGTAGGACC |
| *rel* seq fwd | Amplification of *rel* and flanking sequence for sequencing confirmation | AGGAATAGTATACAAATTAAACTCGC |
| *rel* seq rev | Amplification of *rel* and flanking sequence for sequencing confirmation | GTAGAGTTCTGACCGATACC |
| *relP* upstream fwd | Amplification of *relP* gene half | ATAAGCTTGATATCGATAAGC  TTGATATCGTATCAGCCCAAAGTTCGATAGTG |
| *relP* downstream rev | Amplification of *relP* gene half | ACCGCGGTGGCGGCCTTA  ATTGTTATGTTGTATGTGGGATATTTCTAATTG |
| *relP* upstream rev | Amplification of *relP* gene half lacking nucleotides 450-536 | CATATCCATACCTATAATATAAT  CTTTACGTTTTATCAATTGTACGTCTTC |
| *relP* downstream fwd | Amplification of *relP* gene half lacking nucleotides 450-536 | CGTAAAGATTATATTATAGGTAT  GGATATGTGGGCAAGTTTAG |
| *relP* seq fwd | Amplification of *relP* and flanking sequence for sequencing confirmation | ATCGTACTTTGATAGCGAATCAATTGG |
| *relP* seq rev | Amplification of *relP* and flanking sequence for sequencing confirmation | TCAAAGTCACTCCTTCATTACACG |
| *relQ* upstream fwd | Amplification of *relQ* gene half | ATAAGCTTGATATCGATAAGCTTGATATCGATGC  GTTTATATATTAATGAAATTAAAATTAAAGATGAC |
| *relQ* downstream rev | Amplification of *relQ* gene half | ACCGCGGTGGCGGCCTCA  TCGTTCTTCATCACTTGATATGAAAG |
| *relQ* upstream rev | Amplification of *relQ* gene half lacking nucleotides 343-429 | GAAATTCATTGCTAAACCACTTTCTTTAGTGTTA  CGAATATAATCTC |
| *relQ* downstream fwd | Amplification of *relQ* gene half lacking nucleotides 343-429 | ACTAAAGAAAGTGGTTTAGCAATGAATTTCTGG  GCAAC |
| *relQ* seq fwd | Amplification of *relQ* and flanking sequence for sequencing confirmation | TTGCCATGATATGTATACACCTCG |
| *relQ* seq rev | Amplification of *relQ* and flanking sequence for sequencing confirmation | AGGTATTAAATTAACACTCGGTATTTCTCG |
| *codY* upstream fwd | Amplification of *codY* upstream flanking sequence | ATAAGCTTGATATCGATAAGCTTGATATCGA  TGGATACAGCTGGAATAAGATTAACTCC |
| *codY* upstream rev | Amplification of *codY* upstream flanking sequence | CATGAATTTTTCTCCTTTTGTATATTTTTATA  GAATAAATGC |
| *codY* downstream fwd | Amplification of *codY* downstream flanking sequence | GGAGAAAAATTCATGTAAGTCGATGAGTCT  GGGACATAATTC |
| *codY* downstream rev | Amplification of *codY* downstream flanking sequence | ACCGCGGTGGCGGCCTTATTCTTCAGTTG  CTTCTGTTTCTTCTG |
| *codY* seq fwd | Amplification of *codY* flanking sequence for sequencing confirmation | TATTGGATCAGGAGGCAACTACG |
| *codY* seq rev | Amplification of *codY* flanking sequence for sequencing confirmation | CTGAAATAGTTGCCATTCATTATTCCTCC |
| *lexA* upstream fwd | Amplification of *lexA* gene half | ATAAGCTTGATATCGAACGCTATTTCGC  AAAAATAGGCAAAATAACG |
| *lexA* downstream rev | Amplification of *lexA* gene half | ACCGCGGTGGCGGCCCAGATAAACCAA  AAGATGAGGATATACTTGAACGC |
| *lexA* downstream fwd | Amplification of *lexA* gene half with S130A mutation | CGTAGGCGACGCTATGATTGAG |
| *lexA* upstream rev | Amplification of *lexA* gene half with S130A mutation | AATCATAGCGTCGCCTACGACG |
| *lexA* seq fwd | Amplification of *lexA* and flanking sequence for sequencing confirmation | CTCCTTTGCTTCTTCTTGAGTTAATCC |
| *lexA* seq rev | Amplification of *lexA* and flanking sequence for sequencing confirmation | TATGCACATCAAGGATTAATGAATTCTATTGG |
| pJB185 amp fwd | Amplification of pJB185 backbone | TCTAGAATGACAATGATTACAGATTCATTAGC |
| pJB185 amp fwd | Amplification of pJB185 backbone | AATTCGTAATCATGTCATAGCTGTTTCC |
| pGO1 *rep* fwd | Amplification of pGO1 *rep* promoter region | ACATGATTACGAATTATATAATTGACCTGTGAGGCAAC |
| pGO1 *rep* rev | Amplification of pGO1 *rep* promoter region | CATTGTCATTCTAGACATGATAAAAACTCCTTTAAATGT  ATATTTAAGG |
| pGO1 *trsA* fwd | Amplification of pGO1 *trsA* promoter | TATGACATGATTACGAATTCTGACAAAAAAGTTAAAAAA  GTTTTATAAATTACATGAC |
| pGO1 *trsA* rev | Amplification of pGO1 *trsA* promoter | TCATTGTCATTCTAGACATTTACAACCCACCTTTTCTATT  TGATAAATC |
| pGO1 *aphD* fwd | Amplification of aphD gene fragment using ddPCR | CAAGAGCAATAAGGGCATACC |
| pGO1 *aphD* rev | Amplification of aphD gene fragment using ddPCR | TTCATTGCCTTAACATTTGTGGC |
| *femA* fwd | Amplification of femA gene fragment using ddPCR | CATGGCATTGACCGTTATAATTTC |
| *femA* rev | Amplification of femA gene fragment using ddPCR | CACCAACATATTCAATAATTTCAGCA |

Underlined nucleotides are complementary to the vector and used for In-Fusion cloning

**SUPPLEMENTARY FIGURES**

**n**
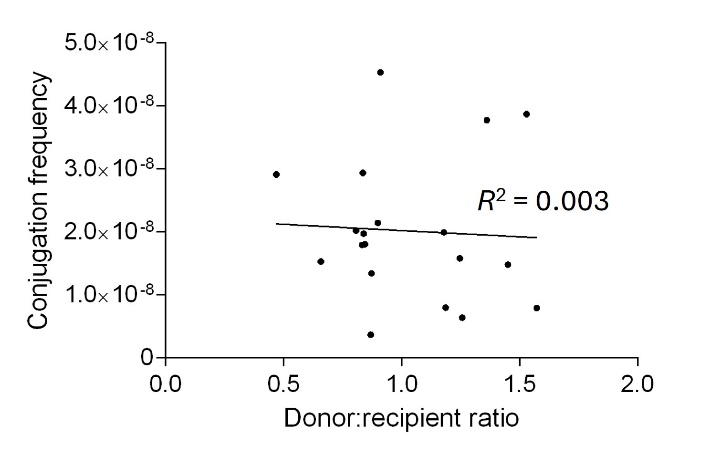

**Figure S1. Effect of donor-to-recipient ratio on conjugation frequency.** Conjugation frequency and donor-to-recipient ratio data for wildtype Newman (pGO1) matings with SH1000-NR were collated from different experiments/days and plotted. A linear regression line was fitted in GraphPad Prism and the goodness of fit (*R*^2^) is shown.

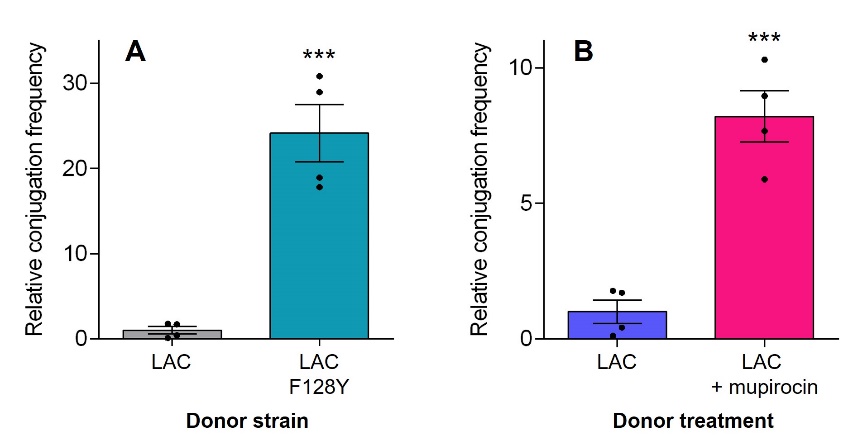

**Figure S2. Rel mutation and mupirocin exposure both increase the rate of pGO1 donation in USA300 LAC.** Filter matings were performed with the donor indicated carrying pGO1, and SH1000-NR as the recipient. For LAC + mupirocin, the donor was exposed to a subinhibitory concentration of mupirocin prior to mating. All conjugation frequencies were calculated as transconjugants per donor and are expressed relative to the LAC mean. Data shown are the mean of four biological replicates; errors bars represent the SEM. Asterisks indicate statistically significant differences between means as determined by a two-tailed *t*-test (*** = *P ≤*0.001).

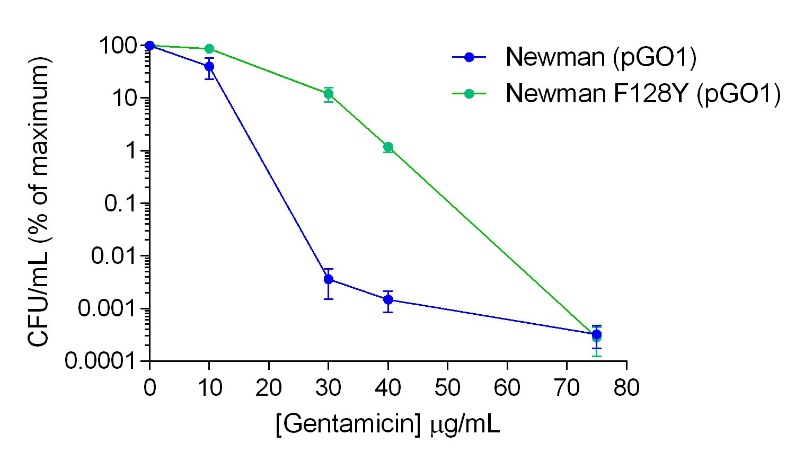

**Figure S3. Population analysis profile of wildtype Newman and the Rel F128Y mutant carrying pGO1 with gentamicin.** Triplicate overnight cultures were diluted, plated on TSA containing increasing concentrations of gentamicin, and the number of colonies counted following incubation. For each replicate culture, the number of CFU/mL at each gentamicin concentration were expressed as a percentage of the maximum CFU/mL determined in the presence of no gentamicin. Error bars represent the SEM.
